## Supplementary material for "Biological networks and GWAS: comparing and combining network methods to understand the genetics of familial breast cancer susceptibility in the GENESIS study": S1 Table

| Network | SNPs | Edges | Subnetworks | $\overline{\text{Betweenness}}$ | $\hat{\mathrm{P}}_{\mathrm{SNP}}$ |
|---|---|---|---|---|---|
| GS | 197 083 | 197 060 | - | $2.03 \times 10^7$ | 0.49 |
| SConES GS | 1 590 | 1 585 | 5 | $2.52 \times 10^7$ | 0.023 |
| GM | 197 083 | 6 442 446 | - | $3.99 \times 10^6$ | 0.49 |
| SConES GM | 1 692 | 177 611 | 5 | $4.40 \times 10^6$ | 0.055 |
| GI | 197 083 | 28 733 720 | - | $1.46 \times 10^6$ | 0.49 |
| SConES GI | 408 | 539 | 5 | $9.33 \times 10^6$ | 0.076 |

$\overline{\text{Betweenness}}$: mean betweenness of the selected SNPs in the corresponding full network.$\hat{\mathrm{P}}_{\mathrm{SNP}}$: median P-value of the selected SNPs.
