## Supplementary figures and images for "Biological networks and GWAS: comparing and combining network methods to understand the genetics of familial breast cancer susceptibility in the GENESIS study"

### S1 Fig

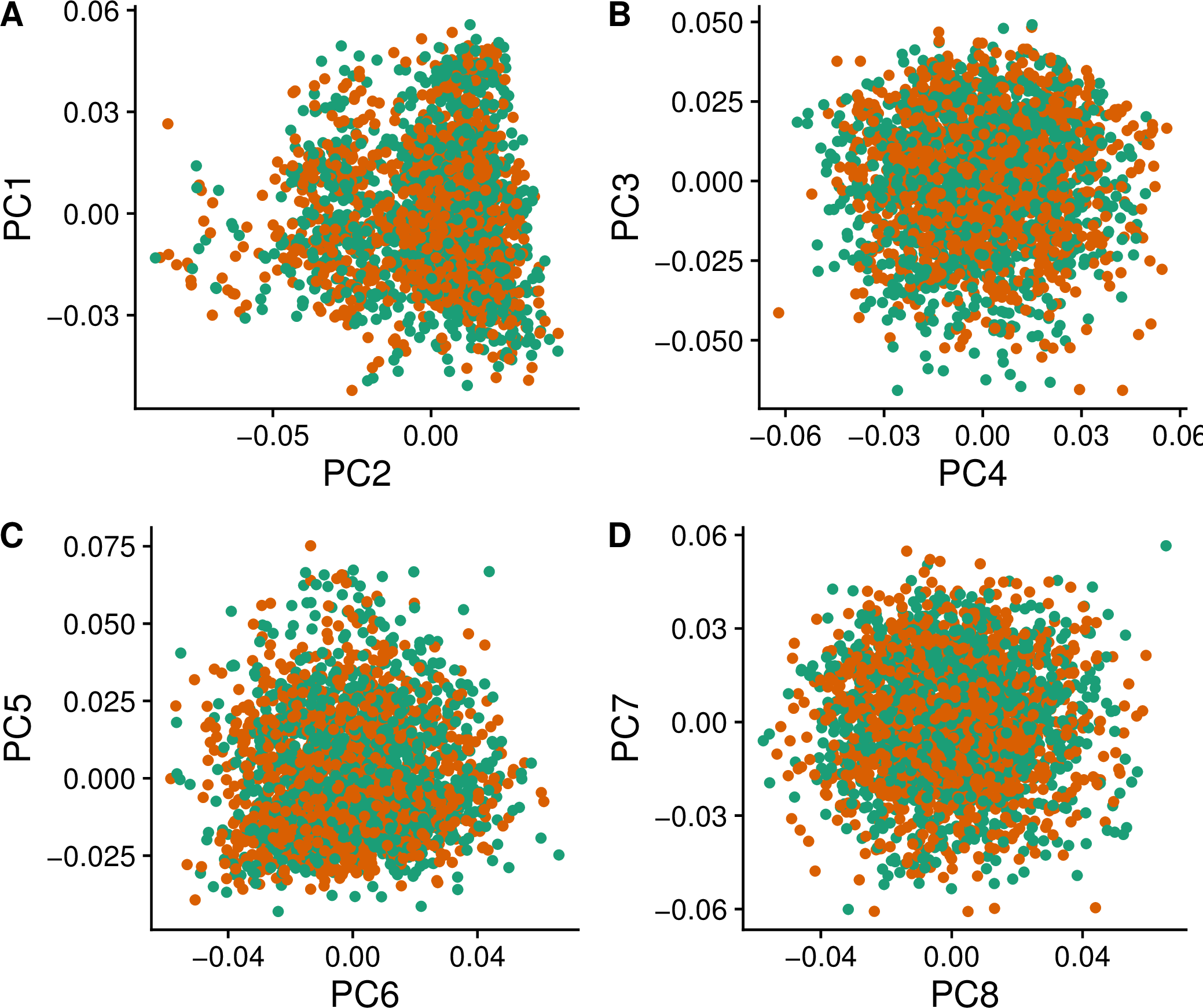

### S2 Fig

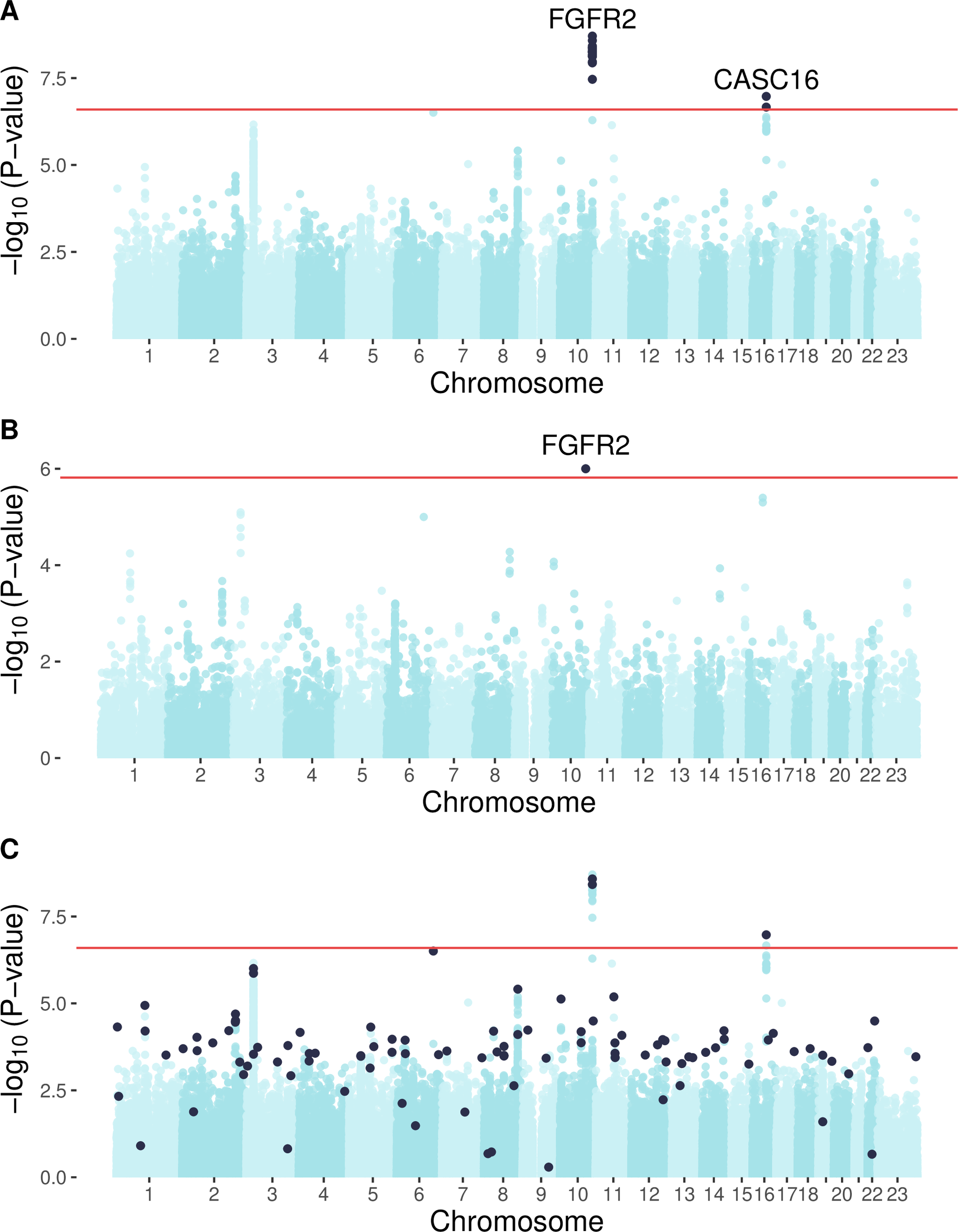

### S3 Fig

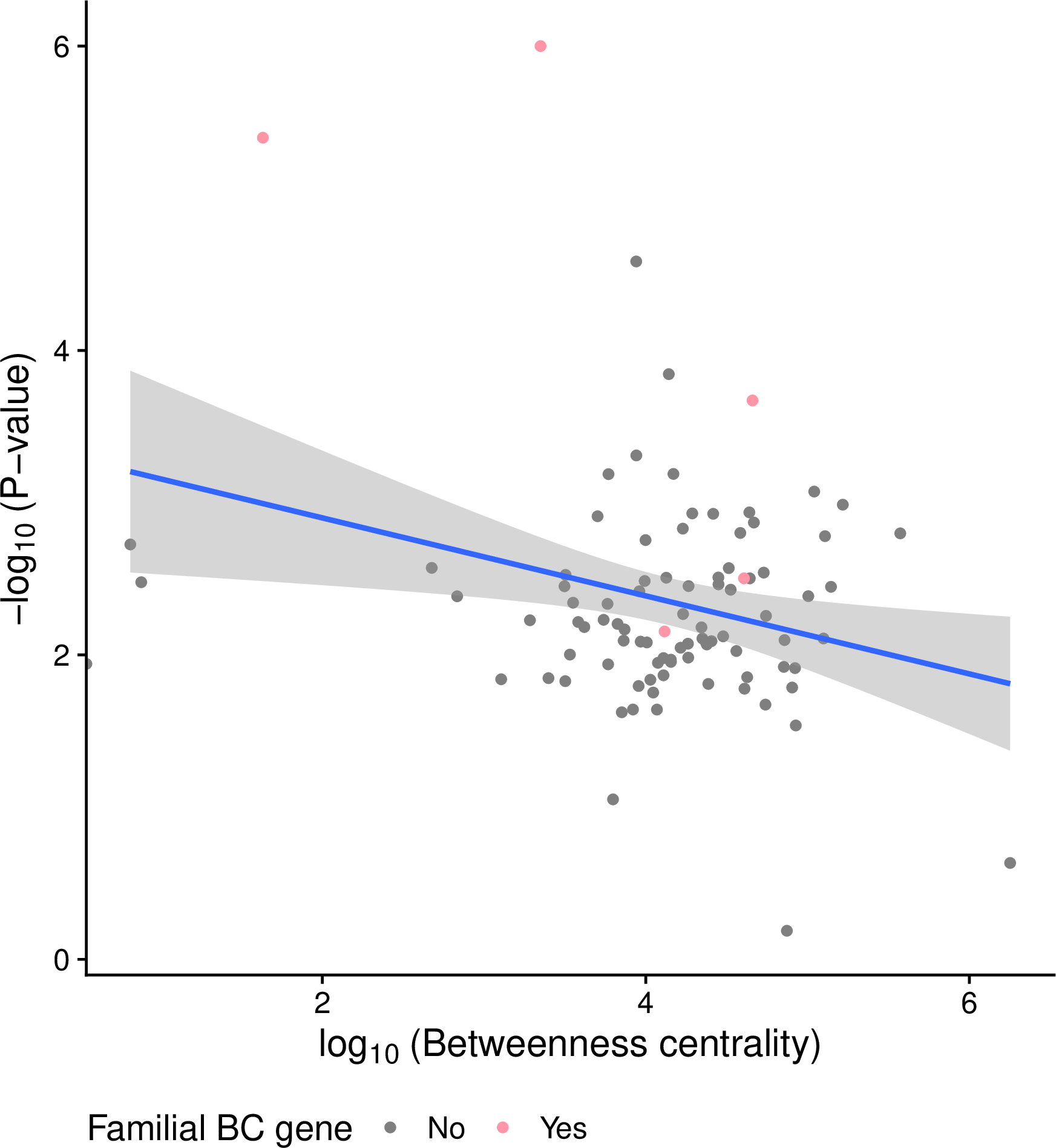

### S4 Fig

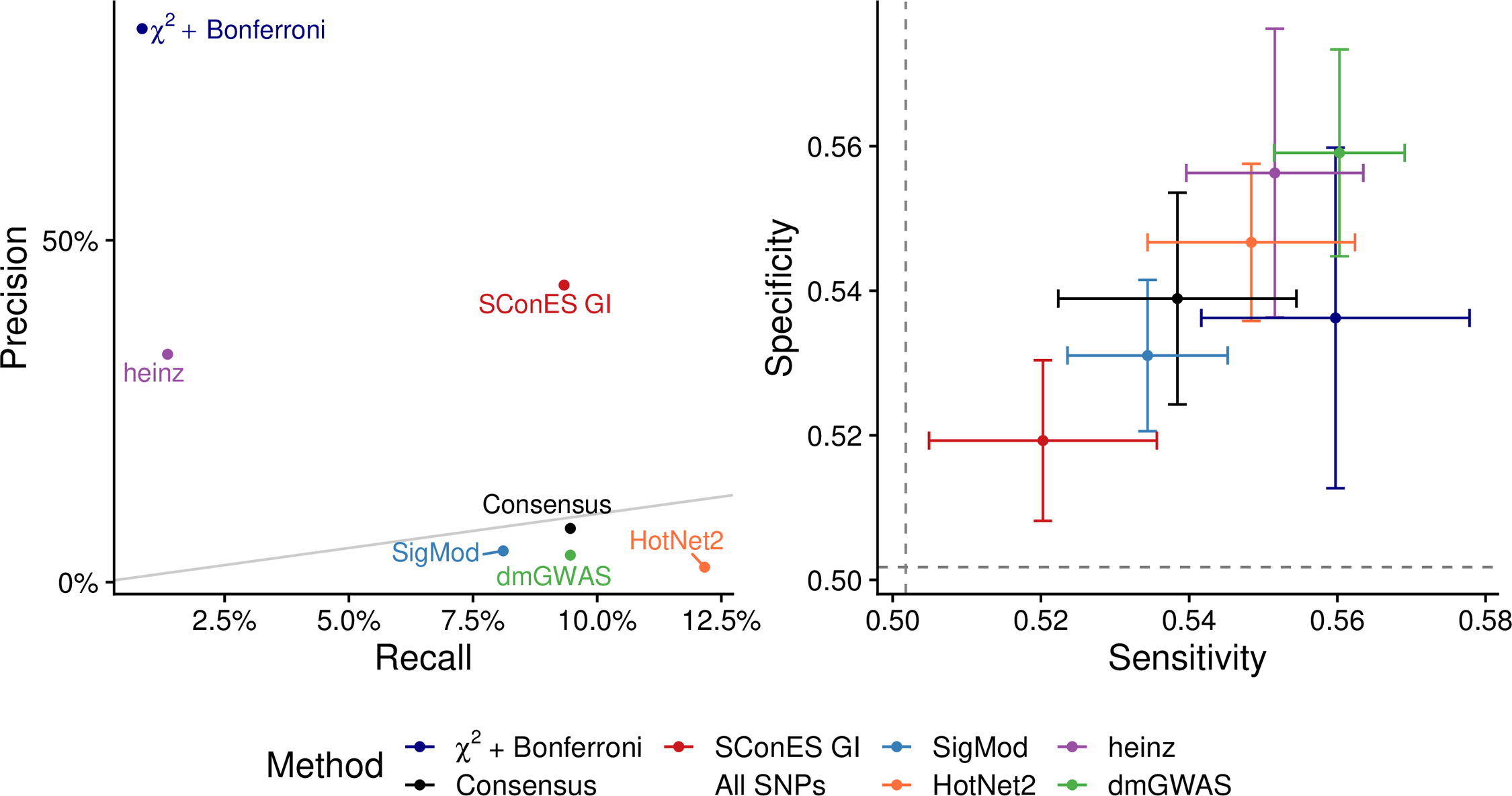

### S5 Fig

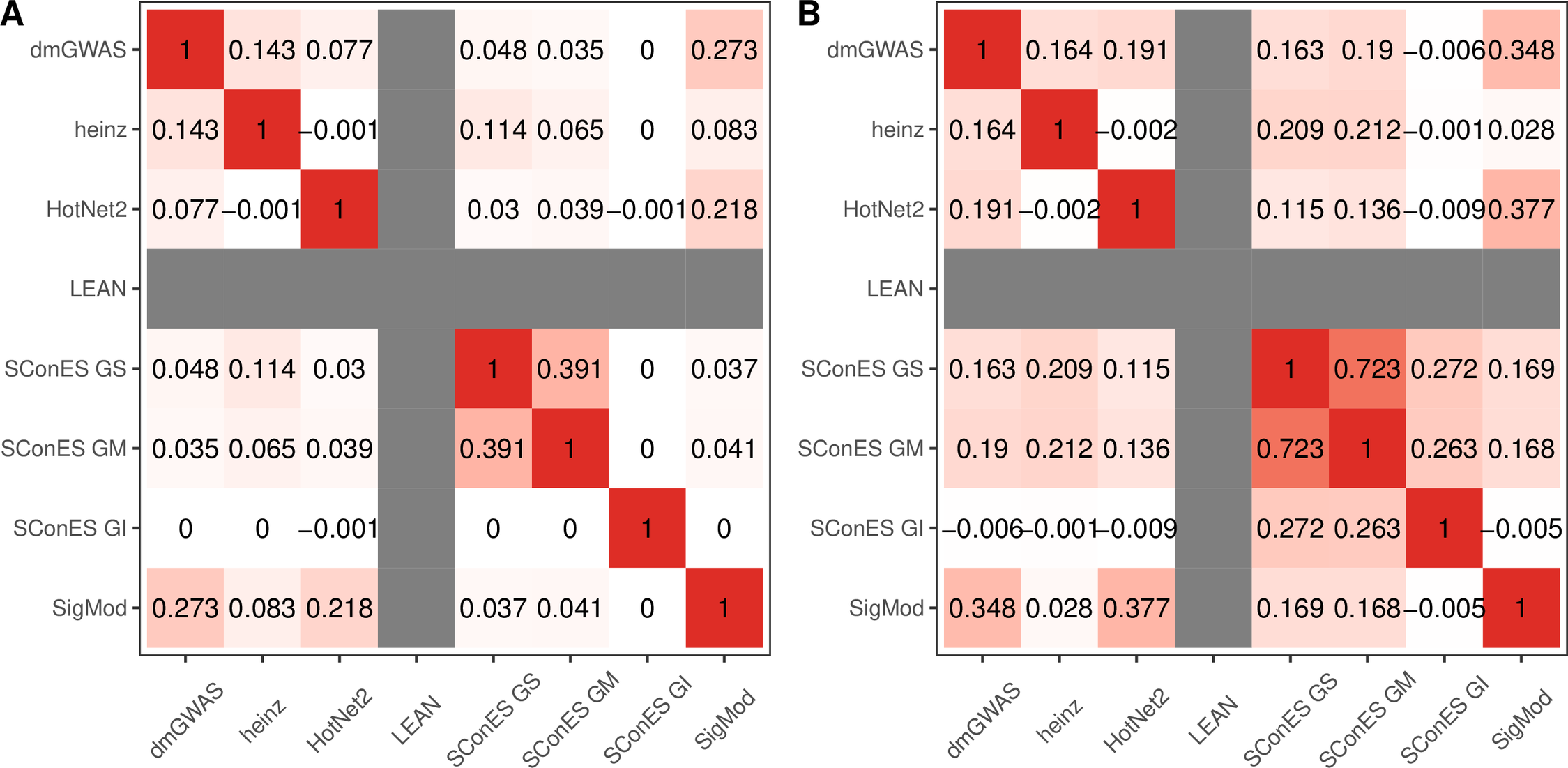

### S6 Fig

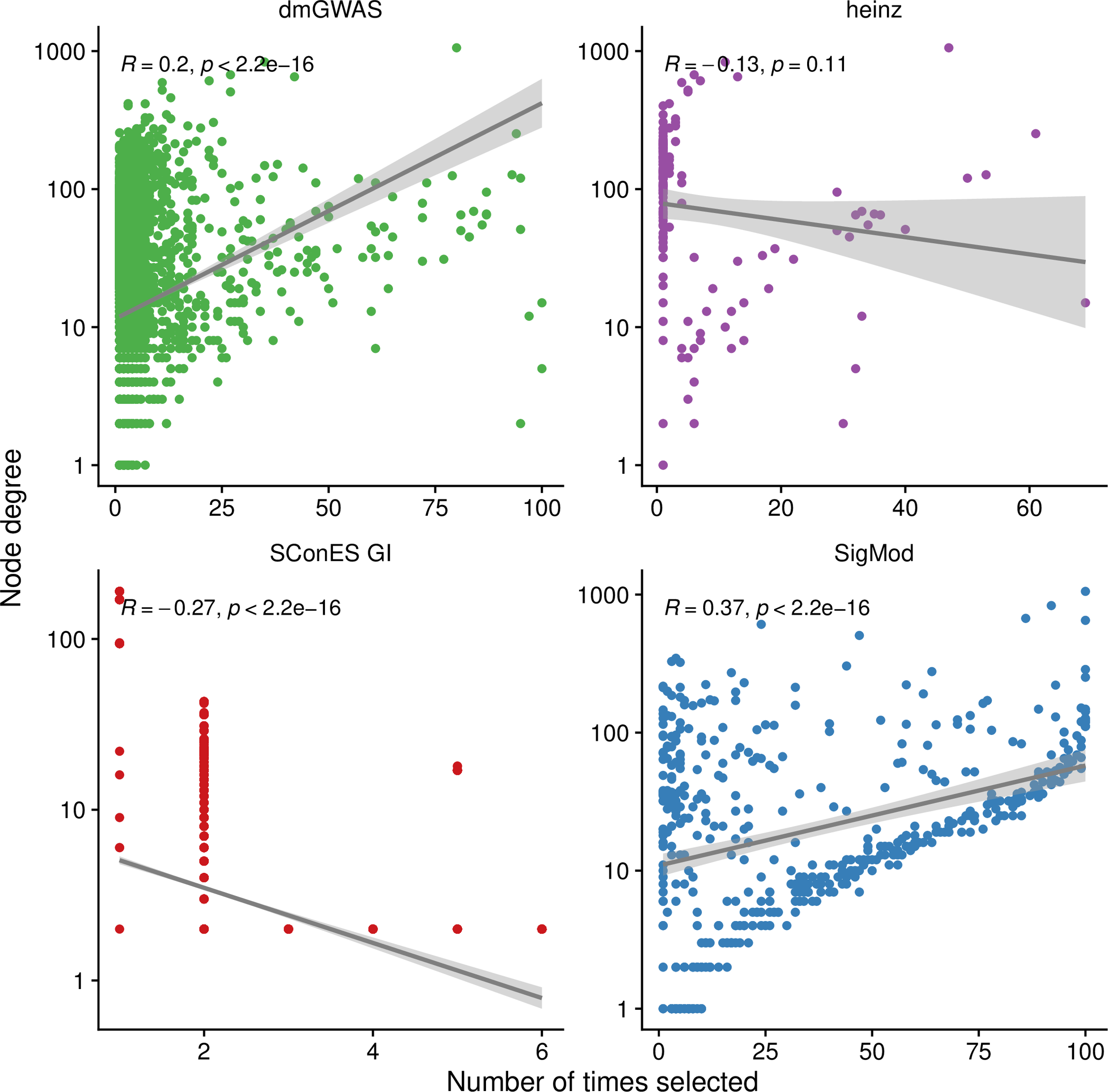

### S7 Fig

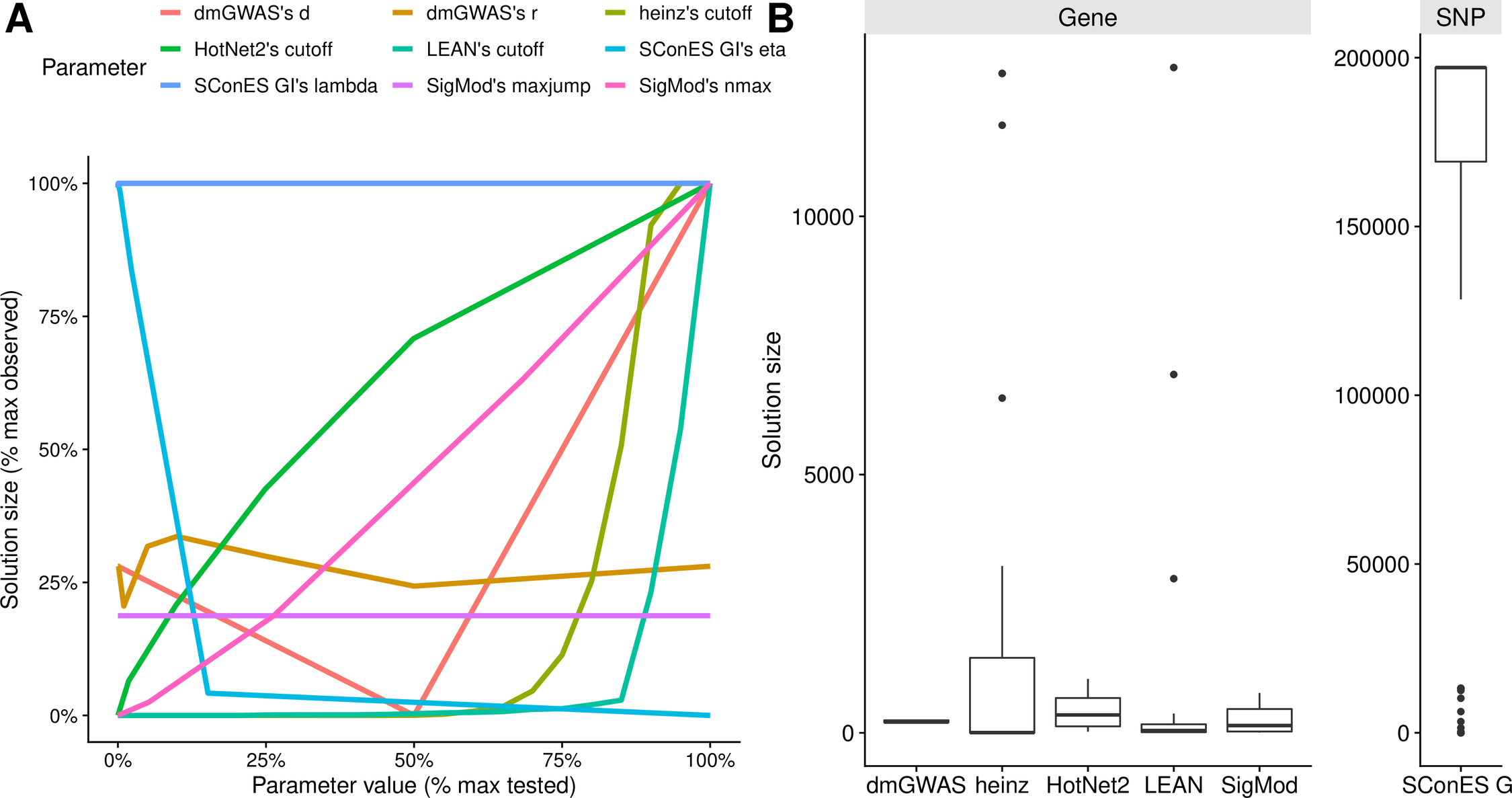

### S8 Fig

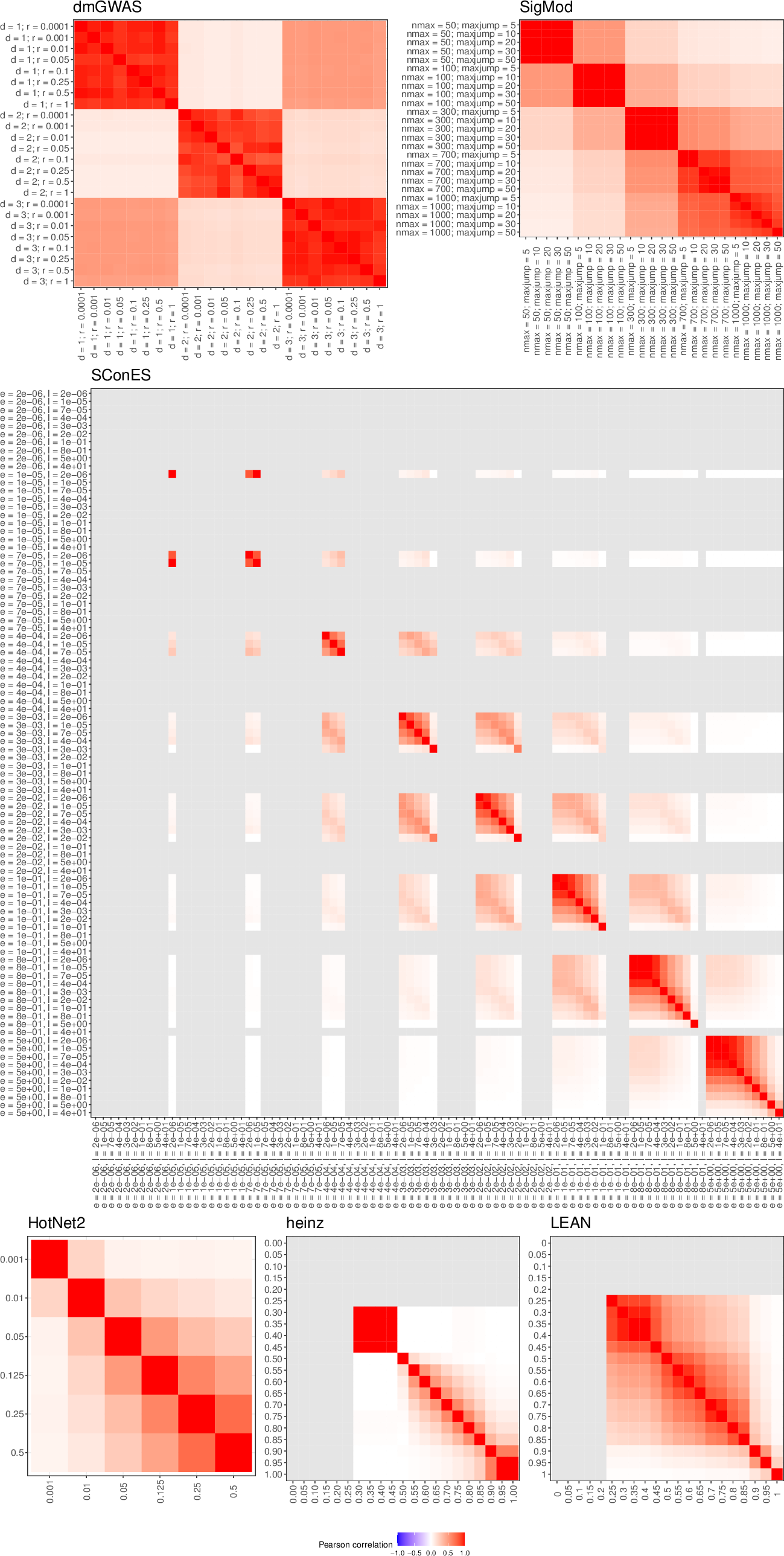

### S9 Fig

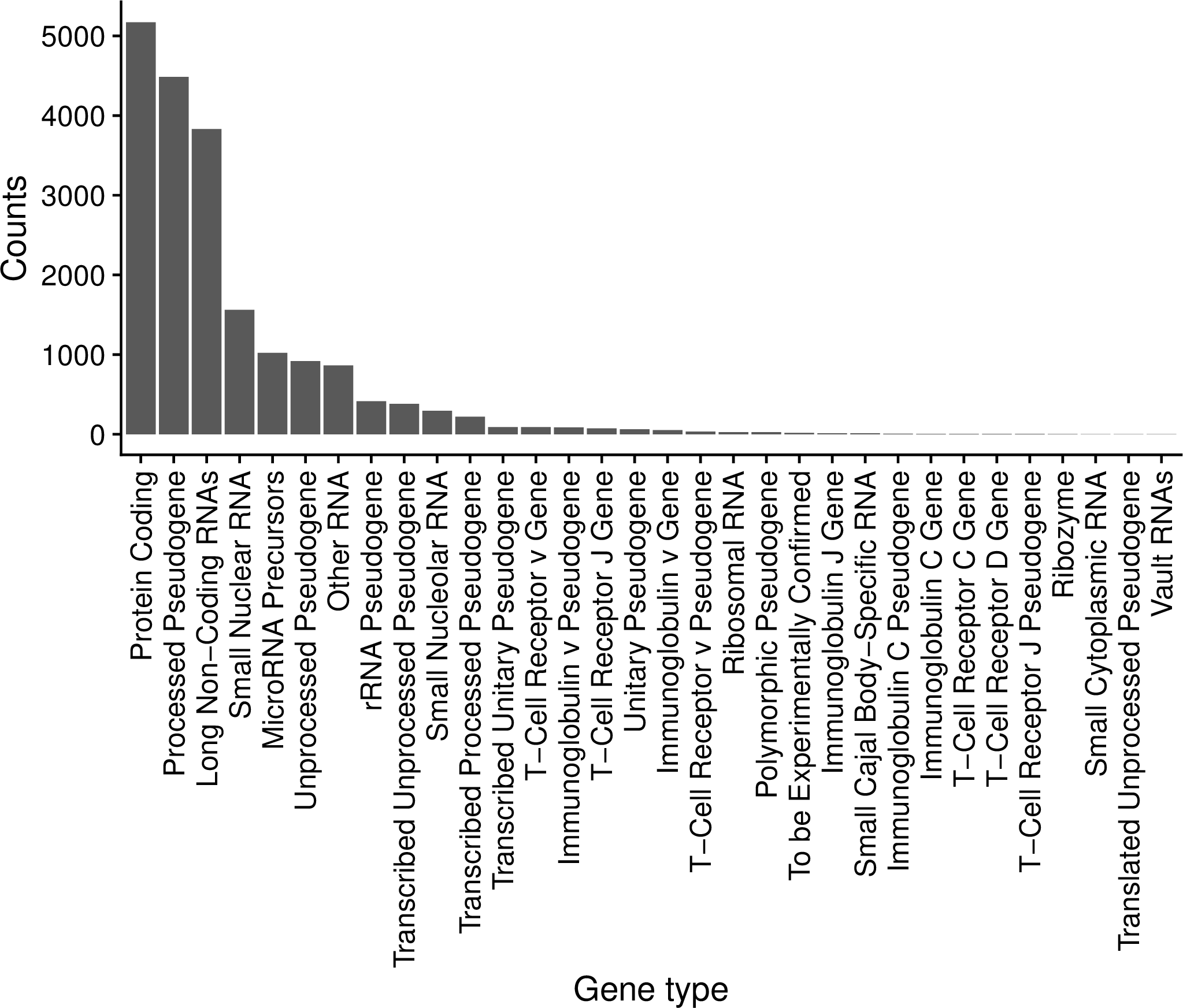

### S10 Fig

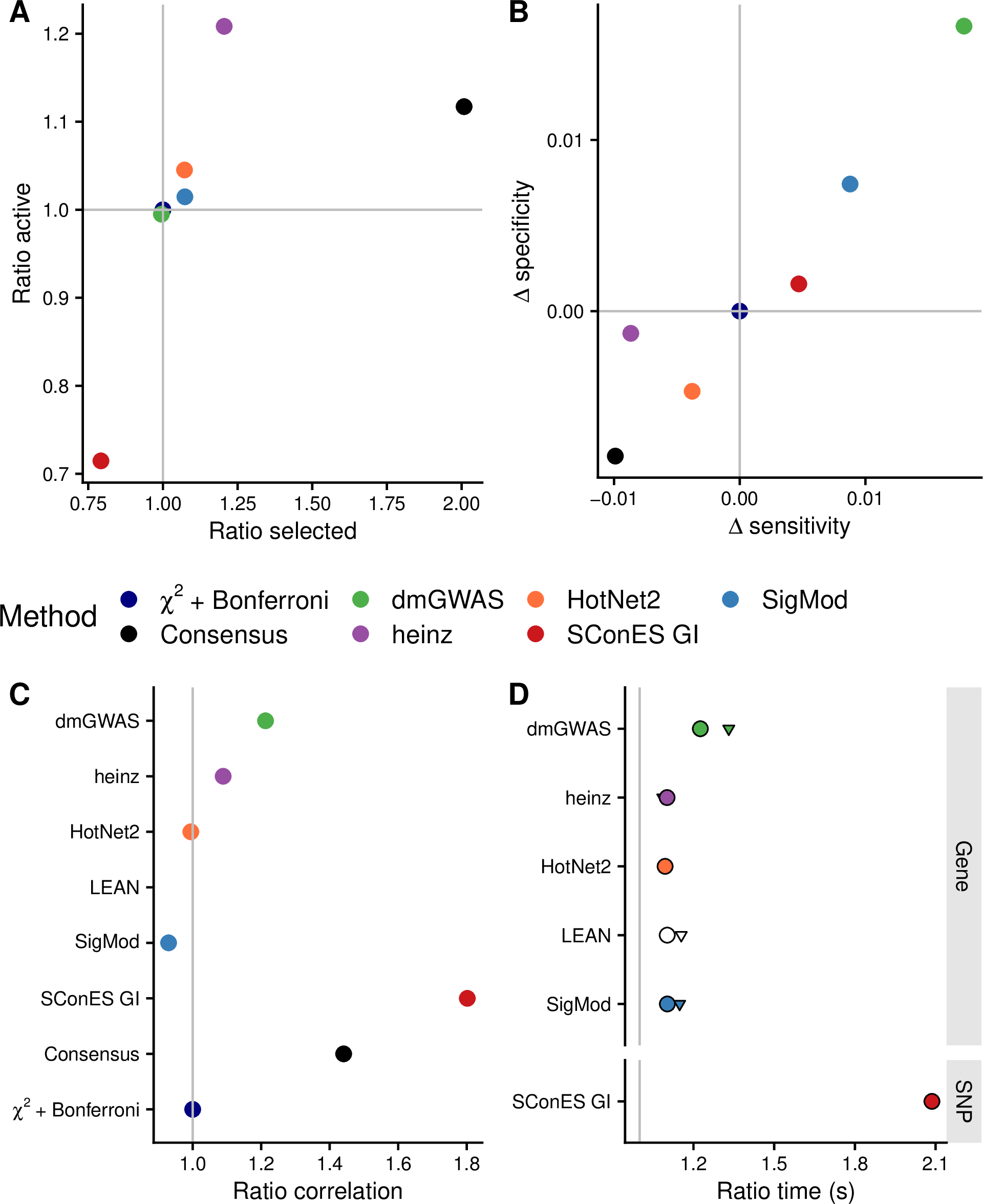

### S11 Fig

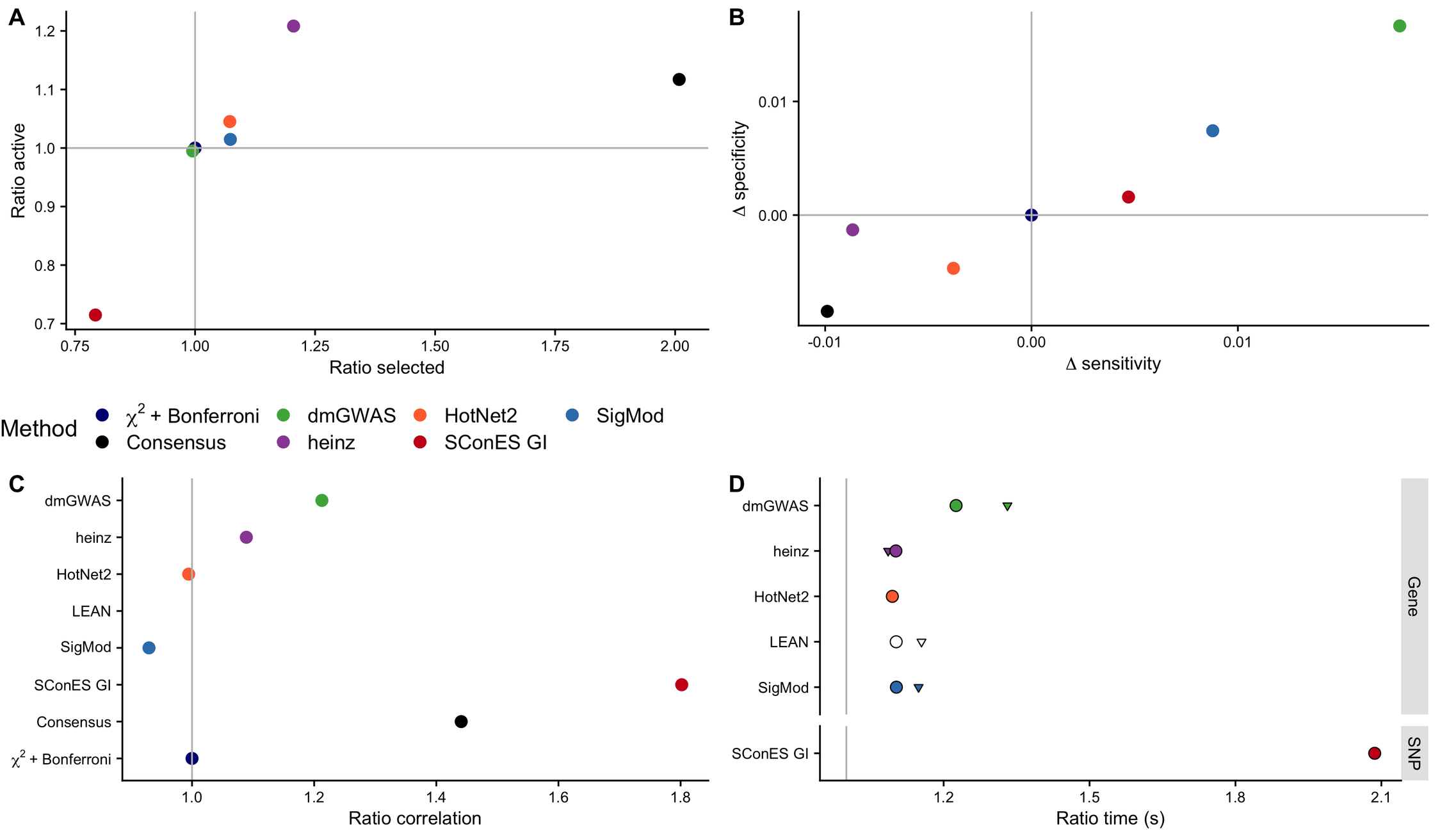
